# How do different functional groups of crop perform in temperate silvoarable agroforestry systems? A case study

**DOI:** 10.1101/2023.02.07.527489

**Authors:** Christina Vaccaro, Johan Six, Christian Schöb

**Affiliations:** Institute of Agricultural Sciences, Department of Environmental Systems Science, ETH Zurich, Zurich, Switzerland; Área de Biodiversidad y Conservación, Universidad Rey Juan Carlos, Móstoles, Spain

**Keywords:** temperate agroforestry, silvoarable, shade, field bean, barley, rapeseed

## Abstract

Agroforestry systems provide a number of ecosystem services and are frequently considered as a promising diversification strategy for more sustainable and climate resilient primary production. Still, less than 1% of the agricultural land in the European Union is silvoarable agroforestry. Most agroforestry field trials compare one crop type with a control in open field with no additional environmental treatments such as nutrient and water availability, thereby limiting our understanding of the ecological processes underlying the potential benefits of agroforestry for food production. The present experimental study addresses three factors (shade, fertilisation, irrigation) on three functionally different crop species (field bean, summer barley, summer rapeseed) and a C_4_-grass (*Echinochloa crus-galli*) in a Swiss agroforestry system. The objective of this study was to assess if and how crop performance (physiological traits, yield) between functional groups varies and if and how shade-induced crop yield reductions diverge between treatment combinations, aiming to provide general functional crop species and management recommendations as a guideline for a successful agroforestry practice in temperate Europe. Summer barley (−44%) and field bean (−38%) showed significant yield declines, similar to summer rapeseed with a significant biomass decline (−35%). Shade significantly increased the occurrence of lodging in barley. Rapeseed in particular performed better when fertilised (+40% biomass). Our results enable to estimate the range of potential yield losses in the competitive zone near mature trees for functionally different crop types and serve as a decision-support for species selection in temperate European agroforestry systems.

## Introduction

Agriculture and climate are interdependent. Both climate change adaptation and mitigation exert mounting strains on agricultural practices. Agroforestry has gained increasing interest due to the strong demand of sustainable and at the same time economic food production (Augère-Granier, 2020). Agroforestry systems (AFS) are long known to deliver a range of ecosystem services such as improved soil fertility, microclimate amelioration, maintenance of air and water quality, weed and pest suppression, biodiversity conservation, erosion control, carbon sequestration and greenhouse gas mitigation (e.g., Kohli *et al*., 2007; Jose, 2009; Torralba *et al*., 2016). Regarding the economic viability of silvoarable AFS, there have always been context-dependent results from various studies, ranging from positive to neutral to negative effects on crop yield. For example, durum wheat (*Triticum turgidum* L. subsp. *durum*) yield was reduced by 13% and 21% in two consecutive years in an AFS with hybrid walnuts (*Juglans nigra* × *Juglans regia* NG23) compared to a pure stand wheat control in Southern France (Dufour *et al*., 2013). Analysing multiple-year (2009–2016) crop yield data of oilseed rape and winter wheat in AFS and control fields, no negative influence on the average long-term crop yields were found in Northern Germany (Swieter *et al*., 2019). In Belgium, two AFS (*Populus* × *canadensis* and various other tree species) were studied to analyse the effects of tree size and age (2-48 years range) on yield and quality of intercrops in a set of 16 arable alley cropping fields (Pardon *et al*., 2018). Near young tree rows, effects on crop yield were limited for all crops, i.e. (forage) maize, potato, winter wheat and winter barley. Near mature trees, yield was substantially decreased, in particular for maize (−65%) and potato (−46%). In contrast, increased winter wheat (*Triticum aestivum* var. Patras) seed yield compared to an open field control was reported within a short rotation alley cropping system with North–South orientated poplar hedgerows in Northern Germany (Kanzler *et al*., 2019). In conclusion, there seems to be no general but highly context-dependent effects of trees on understorey crop yield in silvoarable AFS.

Since trees also provide agricultural products, AFS are potentially more economically productive than monocultures despite reduced understorey crop yields in proximity to trees in mixed systems. For instance, in the Mediterranean climate of Southern France, land equivalent ratios (LER) lay between 1.3 and 1.6 for different combinations of tree-crop systems with an average of 1.2 over the 60-year rotation, despite lower understorey crop yields (Dupraz *et al*., 2018a). Although scientific evidence of the sustainability as well as economic profitability of AFS has amplified rapidly in recent years, mere 9% of the agricultural land in the European Union is agroforestry and 0.39% of the arable lands are silvoarable agroforestry (den Herder *et al*., 2017). Yet the extent of regions and local areas suitable for silvoarable agroforestry is high: Integrating data on soil, climate, topography and land cover in a geographic information system (GIS) to identify regions where *Juglans* spp., *Prunus avium, Populus* spp., *Pinus pinea* and *Quercus ilex* are expected to grow productively and where silvoarable AFS could potentially reduce the risk of soil erosion, nitrate leaching and increase landscape diversity, 56% of the arable land throughout Europe was suitable (Reisner *et al*., 2007). The wide gap between potential and realised silvoarable agroforestry in Europe can be attributed to high implementation costs and a lack of financial incentives, AFS product marketing, education, awareness and field demonstrations (Sollen-Norrlin, Ghaley and Rintoul, 2020), as well as farmers’ doubts about or assumptions of low productivity and profitability (Sereke *et al*., 2016; Rois-Díaz *et al*., 2017). In Switzerland, traditional AFS occupy about 8% of the agricultural lands (Herzog *et al*., 2018; Kay, Jäger and Herzog, 2020) in addition to approximately 250 ha of modern silvoarable agroforestry systems (Kay, Jäger and Herzog, 2020). Meanwhile, the Federal Office for Agriculture (FOAG) has launched an agroforestry project where to date (as of 2022) additional 100 ha of silvoarable AFS were realised (Sonja Kay, personal communication).

An optimisation of silvoarable AFS requires empirical data in the most diverse contexts. Recommendations regarding tree density, arrangement and management have been widely publicised (e.g., within the AGFORWARD innovation and best practice leaflets, see Kanzler *et al*., 2018). Certain functional plant groups have been recommended (such as winter cereals) or found to be inappropriate (such as potato or C_4_ plants like maize) for AFS (e.g., Moreno and Arenas 2017, Pardon *et al*., 2018, Laub *et al*., 2022). However, a comparison of different functional groups of understorey crops under several environmental conditions for silvoarable AFS in temperate climate in an experimental field trial has, to our knowledge, not yet been undertaken.

Tree-crop interactions occur above- and belowground with major consequences for light, water and nutrient availability. Competition for light may outweigh all beneficial effects, such as erosion control, improved microclimate, water availability or soil fertility, leading to substantial understorey crop yield reductions (Ong *et al*., 2015). However, the use of light in AFS can be optimised by a suitable tree-crop-combination (Vandermeer *et al*., 1998; Zhang *et al*., 2018). Indeed, overyielding in mixed cropping systems has often been accredited to increased light use efficiency (Malézieux *et al*., 2009). In AFS positive effects are generally attributed to a complementarity in resource capture by trees and understorey crops, which – apart from light – also include water and nutrients (Cannell, Van Noordwijk and Ong, 1996).

The passive movement of water along a gradient of soil water potential from deeper and wetter soil layers to shallower and drier horizons through tree roots is known as hydraulic lift (Richards and Caldwell, 1987). Trees with deep roots could thus potentially act as “bioirrigators” in AFS (Bayala and Prieto, 2020). However, research has shown the subject of water uptake and competition in AFS to be of great complexity with contrasting results (e.g., Fernández *et al*., 2008; Bayala and Prieto, 2020).

Trees are capable to recycle soil nutrients which would otherwise leach out by their deeper rooting zone (“safety-net hypothesis”). Thus, they increase nutrient use efficiency in the system (Rowe *et al*., 1999; Allen *et al*., 2004). Furthermore, tree litter (leaves, twigs, roots) and the persistence of tree roots stimulate the soil microbiome (Beule *et al*., 2020). In three relatively young (5-to 8-year-old) AFS in Germany, poplar rows increased the abundance of several soil bacterial and fungal groups (Beule *et al*., 2020). In Belgium, the presence of trees in several arable fields increased soil carbon and nutrient concentrations (Pardon *et al*., 2017). Other studies observed nutrient competition between trees and understorey crops (e.g., Gao *et al*., 2013).

The present experimental study addresses the impact of different environmental factors on functionally different understorey crop species in a Swiss AFS to provide field-based evidence which is needed for informed decision-making by farmers, advisory services and policy makers (Sollen-Norrlin, Ghaley and Rintoul, 2020). A cereal, an oilseed crop, a legume and a C_4_ plant were grown in a young AFS with artificial shading and combinations of fertilisation and irrigation treatments, allowing comparisons of the crop’s physiological and economic performances under different environmental conditions. By this, we addressed light, water and nutrient availability in a temperate AFS on four functional crop species groups. We hypothesised that (1) crop performance between the functional groups varies in terms of physiological traits and yield, and that (2) shade-induced crop yield reductions diverge between treatment combinations with the aim to provide general functional crop species and management recommendations as a guideline for a successful agroforestry practice in temperate Europe.

## Materials and Methods

### Study Area

Field experiments were carried out at an organically managed AFS in Windlach, Switzerland. Windlach is located in the Northern part of Kanton Zürich (N 47° 32’ 42.22 E 8° 29’ 0.23) and has a warm and temperate climate (“Cfb” in Köppen and Geiger classification) with a mean annual temperature of 9.6°C and a mean annual precipitation of 1174 mm with highest amounts from May to August (113-121 mm per month) (https://de.climate-data.org/europa/schweiz/zuerich/windlach-121265/, 2022). The farm covers a total area of 23.5 ha and is situated 410 m a.s.l. Apple trees (*Malus domestica*, cv. ‘Heimenhofer’, ‘Schneiderapfel’ and ‘Spartan’) were planted in a density of 37 trees ha^-1^ (10 x 28 m distance within and among tree rows, respectively) on 10.2 ha in West–East orientation in November 2015. Intensive tillage is maintained for weed control. Grown understorey crops are sunflower (*Helianthus annuus*), squash (*Cucurbita* sp.) and winter rye (*Secale cereale*). Artificial pasture and fallow occupy several years in the crop rotation (see SI 1 Tab. 1 in the SI for further details). Soil carbon, nitrogen and total phosphorous content amounted to 1.59% ± 0.28 SD, 0.14% ± 0.03 and 636 ± 150 mg P kg^-1^, respectively, and were lower than C, N and P levels present on average in agricultural soils in Switzerland (C: 3.13%, N: 0.29%, P: 932 mg kg^-1^, source: NABO, personal communication).

### Experimental Design

In proximity (approximately 2 m) to the WE-orientated tree row, alternating shade and control plots were set up on 27 and 28 March 2021 (SI 1 Fig. 1). The shade plots were fabricated by means of artificial shade nets (R.G. Vertrieb, Austria) with 40% opacity to target the desired shading value (i.e. 60% of incident photosynthetically active radiation). This threshold was chosen based on previous experimental studies and models which suggest moderate shade conditions in modern AFS (e.g., Dupraz *et al*., 2018b). Control treatments had no shade net. The nets measured 4 x 4 m, covering 2.5 x 2.5 metres at approximately 1.3 m height. Due to the small subplot sizes for each crop species, shade nets were clamped on all sides (approximately down to 0.4 m distance to the ground on the West, East and South side, and approximately 0.6 m on the North side) to ensure shading at low sun positions (SI 2 Fig. 2a). Field bean (*Vicia faba* ‘Fanfare’), summer barley (*Hordeum vulgare* ‘Atrika’), summer rapeseed (*Brassica napus* ‘Campino’) and common millet (*Panicum miliaceum* ‘Quartet’) were sown below the shade constructions and in the control treatments in 0.85 x 0.85 m wide subplots (0.7225 m^2^) which were marked with bamboo sticks. Spacing between these subplots in a plot amounted to 0.15 m (SI 2 Fig. 2b).

**Fig. 1:**
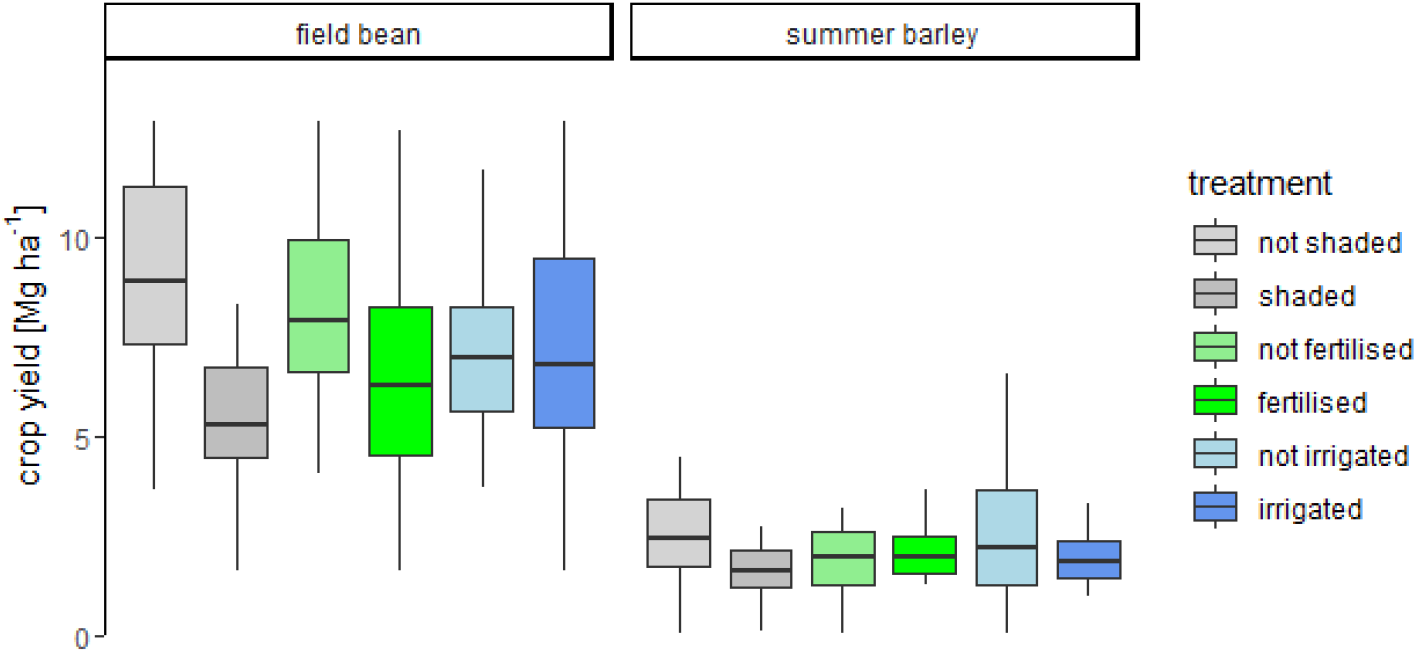
Understorey crop yield under different treatments of shade (not shaded, shaded), fertilisation (not fertilised, fertilised) and irrigation (not irrigated, irrigated) in an agroforestry system in Windlach, Switzerland. The box plots range from the first to the third quartile where the horizontal line shows the median. The vertical lines go from each quartile to the minimum or maximum, respectively.

**Fig. 2:**
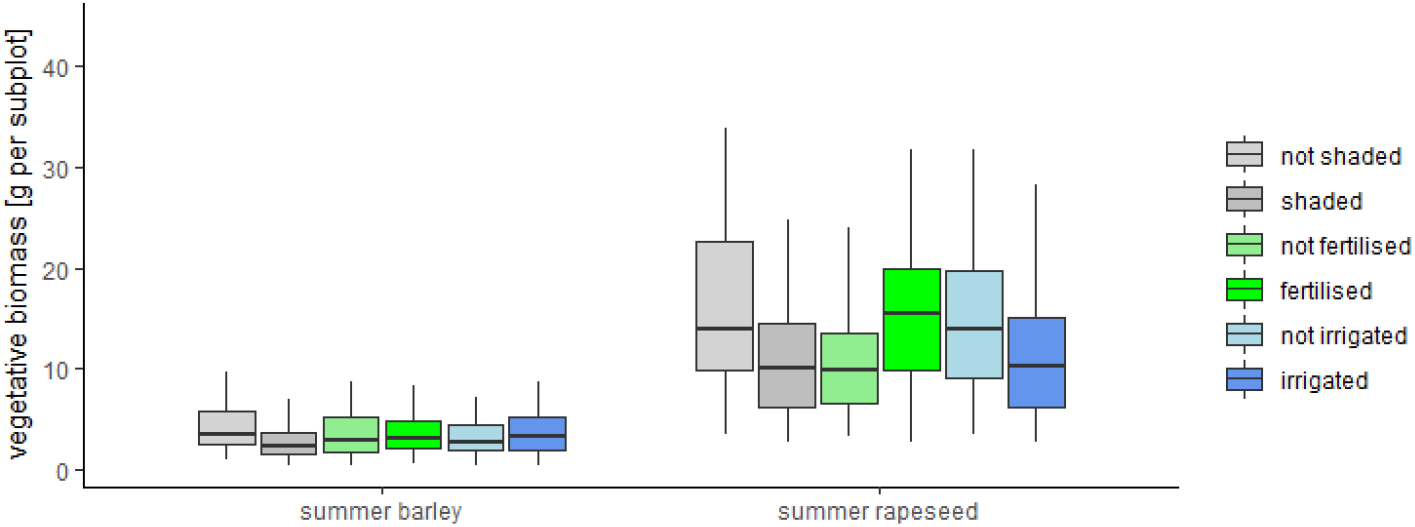
Aboveground vegetative biomass of individual summer barley and summer rapeseed plants under different treatments of shade (not shaded, shaded), fertilisation (not fertilised, fertilised) and irrigation (not irrigated, irrigated) in an agroforestry system in Windlach, Switzerland. The box plots range from the first to the third quartile where the horizontal line shows the median. The vertical lines go from each quartile to the minimum or maximum, respectively.

Soil preparation & sowing: The field was tilled by a rotary tiller (18 March), a plough (25 March) and a harrow (26 March). Field bean and summer barley were sown on 28 March in a density of 61 and approximately 480 seeds per m^2^ (i.e. 44 and 350 per plot), respectively. Summer rapeseed was sown on the 3 April in a density of approximately 140 seeds per m^2^ and common millet was sown on 29 April in a density of approximately 480 seeds per m^2^ (i.e. 100 and 350 per plot, respectively). The seed densities were chosen based on the farmer’s recommendation. Due to absence of germination, common millet was sown a second time 29 May. However, germination failed again. Subsequently, barnyard millet (*Echinochloa crus-galli*) naturally overran the empty millet subplots. As barnyard millet is a C_4_ plant, it was subsequently used as a surrogate of a C_4_ plant for a part of the individual plant trait measurements (plant height, stomatal conductance, chlorophyll content).

Shade (40% opacity) and control (0% opacity) treatments were examined in a full-factorial design with four treatment combinations each: (1) irrigation, (2) fertilisation, (3) fertilisation and irrigation and (4) control, i.e. one replicate consisted of 8 plots (SI 2 Fig. 1). There were four replicates in total.

Fertilisation was carried out according to the fertilisation plan provided by LANDOR fenaco Genossenschaft (SI 2 Tab. 2). The applied organic fertilisers were “Azoplum 13% (Landor)” (N-fertiliser composed of feather-flour), “P 9% Calcophos Landor” (mineral P-fertiliser composed of superphosphate and magnesium oxide), “Calciumschwefel LANDOR” (mineral S-fertiliser composed of Calcium sulfate dehydrate and alpha-cyclodextrin) and “Kalisulfat Streuqualität 50% KaliSOP” (mineral K-fertiliser composed of potassium sulphate).

Irrigation was provided by drip irrigation connected to a total of four 1000 litre water tanks positioned in the tree row. The amount of water was adapted to current weather conditions (SI 2 Fig. 3) and varied between 60-120 litres per plot (i.e. between 10-19 l m^-2^) and watering event (SI 2 Tab. 3). Throughout the growing season only approximately 60 l m^-^2 were irrigated due to the very wet growing season.

**Fig. 3:**
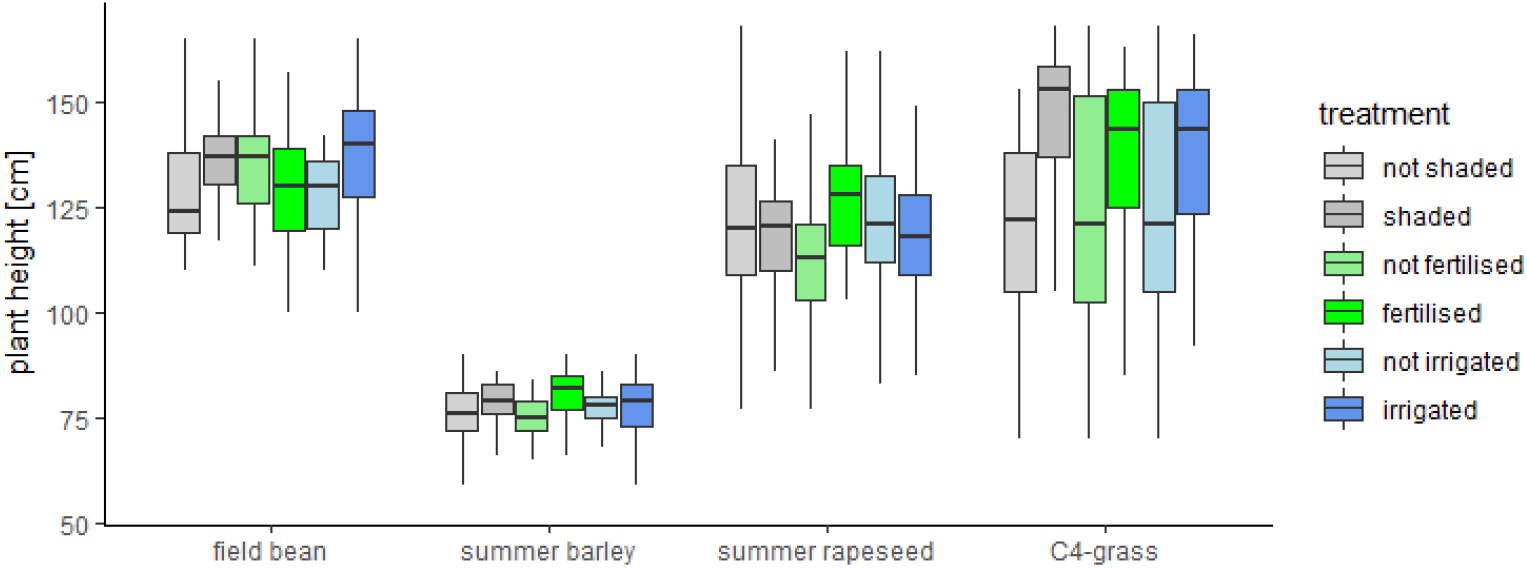
Plant height of field bean, summer barley, summer rapeseed plants and a C_4_-grass (*Echinochloa crus-galli*) under different treatments of shade (not shaded, shaded), fertilisation (not fertilised, fertilised) and irrigation (not irrigated, irrigated) in an agroforestry system in Windlach, Switzerland. The box plots range from the first to the third quartile where the horizontal line shows the median. The vertical lines go from each quartile to the minimum or maximum, respectively.

### Measurements and Sampling

To obtain specific leaf area (cm^2^/g), single leaves of four individuals per subplot were scanned 27 June and then dried for two days at 80°C. Image classification and segmentation was carried out with ilastik (version 1.3.3), pixel count performed with ImageJ (version 1.53n). Subsequently, leaf area was calculated in R (version 3.6.1) by resolution-based conversion of pixel in area. Leaf area (cm_2_) was then divided by leaf dry weight (g).

Leaf chlorophyll content was assessed indirectly by usage of a chlorophyll meter (SPAD-502Plus Konica Minolta®) on 2 and 9-11 July within the four-hour period around noon. The measurement device determines the relative amount of chlorophyll by measuring the absorbance in two wavelength regions where one wavelength corresponds to an absorbance peak of chlorophyll. A numerical SPAD value proportional to the amount of chlorophyll present in the leaf is subsequently calculated from the difference in absorbance. Chlorophyll meter readings show high correlations with extractable leaf chlorophyll and crop N status (Wood, Reeves and Himelrick, 1993), thus allowing comparisons between shade and fertilisation treatments and their controls.

Stomatal conductance was determined with a leaf porometer (SC-1 Leaf Porometer from METER Group ®) on 6 and 9-11 July in the morning and in the afternoon. A leaf porometer measures the actual vapour flux from the leaf through the stomata out to the environment. Stomatal conductance is an indicator of plant water status (Gimenez, Gallardo and Thompson, 2005) and is also affected by light conditions, i.e. it shows a curvilinear relationship with irradiance (Vos and Oyarzún, 1987), thus allowing comparisons between shade and irrigation treatments and their controls.

All subplots were manually harvested when plants reached maturity. For barley, the harvested material (ears with short stalks) was put in labelled paper bags and transported to the ETH Research Station for Plant Sciences in Eschikon (Lindau) where they were threshed with the threshing machine “Saatmeister Allesdrescher K35” (rotational frequency: tumbler: 9, fan: 6). Seeds were weighed and stored in small paper bags in a dry room. For plant trait measurements (plant height, biomass, seed/bean weight, number of seeds/beans), four individuals per species within the area designated for harvest were randomly selected at harvest time. Seed mass was calculated by randomly weighing 10 seeds and dividing the weight by 10. Field bean pods were put in labelled paper bags and dried for four days in two attics. Subsequently, pods of individual samples were opened and the beans were counted and weighed. All beans of one subplot were weighed for total bean weight. For rapeseed, no harvest data could be collected neither on subplot level nor on the individual level due to a flea beetle and rape pollen beetle pest outbreaks, but biomass was collected of four rapeseed plants on the individual level.

### Pests

A heavy infestation of flea beetle (*Phyllotetra* sp.) was observed on 13 of May on five rapeseed subplots, several other rapeseed subplots were also affected. A quick intervention was not possible due to the organic cultivation method. The Research Institute of Organic Agriculture (FiBL) confirmed a difficult year for organic rapeseed production, which was confirmed by the statistical service of the Swiss Farmers’ Union, Agristat, after having collected national yield data. At the beginning of June, first appearances of the rape pollen beetle (*Brassicogethes aeneus*) were noted. After consultation with FiBL, a rock flour suspension was applied twice in June (12 and 19 June). However, the rapeseed subplots could not develop normally due to the two pests, so that measurements and harvest of biomass was only conducted for four of the least affected individuals per subplot.

### Data Analyses

Statistical analyses were carried out with R version 3.6.1. The data was tested for normality and homogeneity of variance by a visual inspection of residuals (normal quantile-quantile plots, standardized residuals versus fits plot) and revision of coefficients of determination (R^2^). Where statistically reasoned, logarithmic or square root transformations were made. On the plot level, lodging and number of plants were tested separately as dependent variable before modelling yield. The number of plants was not significantly influenced by the treatments, thus it was not included as fixed effect in the subsequent analyses. Shade (p < 0.01) significantly increased the occurrence of lodging. Being a consequence of shade, lodging was not included as separate factor but presented separately as an explanatory variable for decreased yields under shade, in particular for barley. Subsequently, statistical modelling was performed with a linear mixed-effects model where species and all treatments (shade, fertilisation, irrigation) and their interactions were tested as fixed effects and replicate and plot as random effects. Species (field bean, summer barley, summer rapeseed, C_4_-grass), shade (0%, 40%), fertilisation (yes, no) and irrigation (yes, no) were factors. The yield on plot level included field bean and barley due to the absence of harvestable rapeseed pods in pest-damaged rapeseed plots and no seeds from inhomogeneous C_4_-grass growth. On the individual level, models followed the same structure, accounting additionally for dependence of individual samples within the same subplots in linear mixed effects models with subplot identification as random term. Apart from seed weight for field bean and barley, rapeseed and C_4_-grass biomass were taken as a proxy for yield given the pest issues with rapeseed that resulted in low and erratic seed yield. Significance of differences between treatments and their interactions was tested by multifactorial ANOVA (type I, sequential sum of squares) and F-tests. For post-hoc analysis, the means of treatment groups were compared with a Tukey test (HSD.test()-function within the R agricolae package, de Mendiburu, 2020) with a significance threshold of α = 0.05.

## Results

### Lodging

Lodging of plants occurred in 11 out of the 128 plots where 10 concerned summer barley and 1 field bean. 10 out of the 11 lodging incidences occurred in shaded plots. Shade (p < 0.01) was significant; species and the interaction of species and shade were highly significant (p < 0.001) for the occurrence of lodging with barley under shaded conditions being most prone to lodging.

### Yield

Species (p < 0.001), shade (p = 0.001), fertilisation (p = 0.025) and the interaction of species and shade (p = 0.049) significantly affected understorey crop seed yield at the plot level (field bean and barley only). Across all fertilisation and irrigation treatments, field bean and barley yield in non-shaded plots amounted to 650 ± 187 SD Mg ha^-1^ and 205 ± 137 Mg ha^-1^ compared to 401 ± 133 Mg ha^-1^ (−38 %) and 115 ± 52 Mg ha^-1^ (−44 %) in shaded plots, respectively (Fig. 1). Across all shade and irrigation treatments, field bean and barley yield in non-fertilised plots amounted to 594 ± 188 Mg ha^-1^ and 188 ± 139 Mg ha^-1^ compared to 461 ± 204 Mg ha^-1^ (−22 %) and 170 ± 72 Mg ha^-1^ (−10 %) in fertilised plots, respectively. Both the shade- and fertiliser-induced yield reductions were significant for field bean but not for barley in the post-hoc Tukey test at the plot level.

At the individual level, species (p < 0.001), shade (p < 0.001) and the interaction of species and shade (p < 0.002) significantly affected bean and seed yield of field bean and summer barley, respectively. Again, yield decrease was significant for field bean but not for barley based on the post-hoc test. Shade significantly decreased total seed number of barley from on average 60 (± 28) to 39 (± 23) seeds per individual (−35%, p < 0.001), total seed weight from 3.3 (± 1.8) to 2.0 (± 1.3) g per individual (p < 0.001), seed mass from on average 0.055 (± 0.009) to 0.050 (± 0.001) g per individual (p <0.05) and the numbers of tillers from on average 9 (± 3) to 7 (± 2) (p < 0.001), respectively. Shade significantly decreased the number of field bean pods from on average 15 (± 6) to 10 (± 3) per individual (−33%, p < 0.001) and the number of beans from on average 40 (± 16) to 31 (± 10) per individual (−22%, p < 0.05). In addition, the interaction of shade and irrigation was significant for pod number (p < 0.01) where in shaded plots irrigation decreased pod number while it was increased in unshaded plots. Likewise, irrigation decreased bean number in shaded plots but increased bean number in unshaded plots, though this was not significant. Shade decreased total bean weight from on average 20 (± 8) to 14 (± 5) g per individual (−30%, p < 0.001). In addition, in the post-hoc test fertilisation significantly reduced total bean weight from 18 (± 8) to 15 (± 6) g per individual (−17%).

Aboveground biomass was significantly dependent on species (p < 0.001), shade (p < 0.001), fertilisation (p = 0.03) and the interaction of species and irrigation (p < 0.05). The reduction of biomass under shade was from 16.4 ± 8.8 g to 10.7 ± 5.3 g (−35%) for rapeseed; and from 4.3 ± 2.3 g to 2.8 ± 1.8 g (−58%) for barley (Fig. 2). Fertilisation increased rapeseed biomass from 11.3 ± 6.8 g to 15.8 ± 8.1g (+40%). Barley biomass was nearly the same (3.5 ± 2.2 g in fertilised, 3.7 ± 2.3 g in non-fertilised subplots). Species and fertilisation interaction was marginally significant (p = 0.06). Irrigation reduced rapeseed biomass (from 15.2 ± 8.1 g to 11.9 ± 7.1 g, −22%) but increased barley biomass (from 3.4 ± 2.0 g to 3.8 ± 2.3 g, +12%).

### Plant Height

Species (p < 0.001), shade (p < 0.05), fertilisation (p < 0.01), irrigation (p < 0.05), the interactions of species and shade (p < 0.05), species and fertilisation (p < 0.01) and species and irrigation (p = 0.05) significantly affected plant height. Among all four functional groups, shade, fertilisation and irrigation increased plant height (Fig. 3). Looking at the interaction of species and shade, the increase in plant height was significant for the C_4_-grass (142 ± 24 cm in comparison to 120 ± 22 cm, +18%). Fertilisation-induced increase in plant height was significant for the C_4_-grass (137 ± 21 cm compared to 124 ± 28 cm, +10%) and rapeseed (127 ± 15 cm compared to 113 ± 16 cm, +12%). Equally, irrigation-induced increase in plant height was significant for the C_4_-grass (137 ± 22 cm compared to 124 ± 27 cm, +10%) and field bean (138 ± 14 cm and 128 ± 9 cm, +8%).

### Chlorophyll Content

Date (p < 0.0001), species (p < 0.0001) and the interaction of species and shade (p < 0.05) had a significant effect on leaf chlorophyll content. For field bean, leaf chlorophyll content was significantly higher in unshaded (54 ± 5) than in shaded controls (51 ± 5) in the post-hoc Tukey test. For barley, rapeseed and C_4_-grass no significant differences were found among shade treatments.

### Stomatal Conductance

Date and species had a significant effect on overall, minimal and maximal stomatal conductance (SC) (p < 0.0001). The interaction of species and shade was marginally significant for minimal SC (p = 0.089). Results of the individual analyses of the effects of shade, fertilisation and irrigation are: (1) significant effects of shade (p < 0.01), fertilisation (p < 0.05) and a marginally significant interaction of shade and fertilisation (p = 0.056) for minimal SC of rapeseed and (2) a marginally significant effect of fertilisation (p = 0.078) for maximal SC of rapeseed. Minimal SC of rapeseed amounted to 364 ± 142 mmol m^-2^ s^-1^ in unshaded plots compared to 440 ± 132 mmol m^-2^ s^-1^ under shade (+21%) and to 390 ± 147 mmol m^-2^ s^-1^ in fertilised plots compared to 412 ± 138 mmol m^-2^ s^-1^ in unfertilised plots (+6%). Shaded, fertilised rapeseed had the highest minimal SC (453 ± 127 mmol m^-2^ s^-1^), unshaded, fertilised rapeseed the lowest minimal SC (328 ± 141 mmol m^-2^ s^-1^, −28%). The difference between shaded, unfertilised rapeseed (429 ± 139 mmol m^-2^ s^-1^) to unshaded, unfertilised rapeseed (395 ± 139 mmol m^-2^ s^-1^) was less pronounced (−8%).

Minimal SC under shade was higher for barley and the C_4_-grass but lower for field bean compared to unshaded plots. Maximal SC was higher under shade for field bean and the C_4_-grass but lower for barley compared to unshaded plots. However, these findings were not significant.

### Specific Leaf Area

Species (p < 0.0001), shade (p < 0.0001), the interaction of species and fertilisation (p < 0.0001) and the interaction of species and shade (p = 0.0001) had a significant effect on specific leaf area (SLA). The three-way interaction of species, shade and fertilisation was marginally significant (p = 0.087). Shade significantly increased SLA across all species. In the post-hoc Tukey test, SLA of field bean, barley and rapeseed was significantly higher in shaded plots (94 ± 9, 87 ± 13 and 69 ± 8 cm^2^ g^-1^, respectively) than in controls (84 ± 11, 77 ± 13 and 53 ± 8 cm^2^ g^-1^, respectively). The shade-induced increase in SLA amounted to 12%, 13% and 30% for field bean, barley and rapeseed, respectively. In the post-hoc Tukey test, SLA of field bean and barley was not significantly different in fertilised plots compared to non-fertilised controls. SLA of rapeseed was significantly lower in fertilised plots (56 ± 11 cm^2^ g^-1^) than in unfertilised plots (65 ± 10 cm^2^ g^-1^, +16 %). Concerning the triple interaction between species, shade and fertilisation: Unshaded rapeseed had a significantly lower SLA when fertilised (49 ± 10 cm^2^ g^-1^ compared to 59 ± 6 cm^2^ g^-1^ when unfertilised). Shaded rapeseed had a lower SLA too when fertilised, but this difference was not significant (67 ± 7 cm^2^ g^-1^ compared to 70 ± 9 cm^2^ g^-1^ when unfertilised).

## Discussion

Despite well-acknowledged response differences of crop species to shade, most studies test only one functional crop type in the same location (Laub *et al*., 2022). The present study investigated three common crops and a C_4_-grass to compare the performance of a cereal, an oilseed crop, a legume and a C_4_ plant in a temperate AFS with controlled environmental treatments of shade, fertilisation and irrigation, addressing light, water and nutrient availability at the same time. Our results demonstrate that interactions between species and environmental factors are significant. Despite of the fact that barley and field bean are considered shade tolerant crops, they showed a similar shade-induced yield decrease as the biomass decrease of the supposedly less shade tolerant rapeseed. For rapeseed, fertilisation clearly had the biggest impact on biomass production and plant height. Irrigation-induced increase in plant height was significant for field bean and the C_4_-grass while it was not for barley and rapeseed.

The acute and severe pest damage, the germination failure for common millet and the unforeseen rainfall quantities impaired the initial expected range of contrasts in functional groups and environmental conditions to some extent. However, the presented results still hold explanatory power. The interaction of species and shade significantly influenced crop yield reduction, i.e. field bean showed a more consistent yield reduction than summer barley at the subplot level, though barley apparently had a stronger decrease (−44% compared to −38% in field bean). Shade-induced yield reduction in barley was in the range predicted by Laub *et al*. (2022) in their meta-regressions for C_3_ cereals, which amounted to −38% under 40% shade. In our study, yield variability was high and standard deviation amounted to 67% and 45% in non-shaded and shaded plots, respectively. Possibly bird grain foraging was reduced under the shade constructions. All yield-related traits (total seed number, seeds per individual, total seed weight, seed mass, number of tillers) and biomass of barley were significantly reduced under shade. Lodging is known to severely decrease cereal yields and is most often attributed to an increase in plant height (Shashidharaiah, 2008). In the AFS in Windlach, shade increased plant height and lodging occurred in 50% of the shaded plots (8 out of 16). Decreased biomass and increased plant height and SLA are classical traits of shade avoidance response (Carriedo, Maloof and Brady, 2016). However, it can be expected that winter barley would perform better in an AFS with deciduous trees as it could benefit from full sun conditions before tree leaf emergence (Nerlich, Graeff-Hönninger and Claupein, 2013).

Field bean showed similar yield drops to barley, with reduced numbers of pods and beans and a decreased average bean weight. Yield reduction of −38% was lower than predicted by the meta-regressions form Laub *et al*. (2022) for grain legumes, which amount to −50% for grain legumes under 40% shade. The authors classified grain legumes as shade susceptible with a more than proportional yield decrease in response to limited solar radiation. Other studies concluded that legumes and legume varieties are relatively shade-tolerant (Hadi *et al*., 2006; Nasrullahzadeh *et al*., 2007). In our study, field bean showed a similar degree of the shade avoidance response as barley, with increased SLA and plant height. In addition, a significant drop in chlorophyll content was observed with shade, as observed in other studies (e.g., Mauro *et al*., 2011; Muhidin *et al*., 2018).

Experimental studies with rapeseed in AFS are scarce. In an alley cropping AFS in northern Germany, rapeseed and winter wheat yields in two years were on average 23% and 45%, respectively, lower at 1 m distance from the tree strip edges compared to the middle of the crop alley (Swieter *et al*., 2019). The stronger reduction for the cereal than the oilseed is in alignment with our findings where shade-induced biomass reduction was stronger for barley compared to rapeseed. Rapeseed showed a particularly strong response (+30% in SLA) to shade as well as a significant interaction of shade and fertilisation, underlining the below-ground competitiveness of rapeseed. Fertilisation was a significant factor for several physiological traits, in particular for rapeseed. Biomass reduction caused by the absence of fertilisation was stronger for rapeseed than barley, corresponding to an observation in an AFS in Southern China where Cao, Kimmins and Wang (2012) found the strongest root competition by rapeseed. Fertilisation also significantly increased plant height in rapeseed. Plant height, though commonly an indicator for growth under shade, can also be increased by fertilisation (Bybordi and Ebrahimian, 2013).

The C_4_-grass was expected to display a distinct yield drop due to its photosynthetic metabolism (e.g., Taiz *et al*., 2015; Gao *et al*., 2020), though shade-tolerant C_4_-grasses exist (Horton and Neufeld, 1998). In this study, not all physiological traits were assessed for the C_4_-grass as a substitute for common millet, but with regard to plant height it is clear that shade had the biggest impact on growth; followed by fertilisation and irrigation which had a similar high effect on growth. In Belgium, yield decrease near mature trees in two AFS was substantial for maize (another C_4_ plant) with −65%. Due to the strong reaction to shade in plant height, it may be assumed that the C_4_-grass (*Echinochloa crus-galli*) was also strongly affected yield-wise.

To summarize: We hypothesised that (1) crop performance between the functional groups varies in terms of physiological traits and yield. In terms of yield, field bean and barley most strongly and negatively reacted to shade. Rapeseed (biomass) negatively responded to shade and in addition positively to fertilisation. The intensity of yield responses was generally in line with the physiological responses of the species to the treatments. We further hypothesised that shade-induced crop yield reductions diverge between treatment combinations. In the instance of rapeseed this was true for SLA where unshaded rapeseed had a significantly lower SLA when fertilised. Lastly, it was hypothesised that general functional crop species recommendations can serve as a guideline for a successful agroforestry practice. Since the cultivation of millet did not succeed, the biggest functional contrast is missing. The shade tolerance and manifold proven suitability of C_3_ cereals in agroforestry (e.g., Dupraz *et al*., 2018a; Pardon *et al*., 2018; Arenas-Corraliza *et al*., 2019; Kanzler *et al*., 2019) is not supported by our data. However, the experimental setup required a summer cereal which falls short of the else provided temporal advantages of early development before leaf emergence of trees. As a recommendation, a greater plant height in response to shading certainly leads to more frequent lodging and should therefore be considered when selecting cereal varieties for AFS. Field bean as a grain legume proved to show a significant yield decrease, but less than generally expected for this functional group (Laub *et al*., 2022).

Lastly, it must be emphasized that in any AFS, including those with mature trees, the actual competition zone beside tree rows is limited to a few metres and followed by a zone of maximum protection (e.g., Pardon *et al*., 2018, Kanzler *et al*., 2019). In the short rotation alley cropping system, for example, the average winter wheat yields decreased by only 1% within 3 m distance of the North-South orientated hedgerow (Kanzler *et al*., 2019). In their study, the area of shelter protection reached from 3 to 24 m from the hedgerow with increased crop yields. Our findings represent potential yield decreases in the immediate vicinity and Northern side of tree rows and do only deliver an estimate of the range of maximum yield declines in the competition zone.

## Conclusion

In this experimental study combinations of shade, nutrient and water availability were tested on four crop species of different functional groups in a temperate silvoarable AFS. The grain legume (field bean) showed the strongest shade-induced yield decline, followed by the summer cereal (barley) and oil crop (rapeseed). Shade significantly increased the occurrence of lodging in barley. Rapeseed performed better when fertilised. Our results enable to estimate the range of potential yield losses in the competitive zone near mature trees for different crop types and serve as an aid to decision-making for species selection in AFS.

## Declarations

### Funding

The authors did not receive specific funding for the submitted work.

### Conflicts of interest

The authors have no conflicts of interest to declare that are relevant to the content of this article.

## Acknowledgments

We thank Adrian Meierhofer for his permission to install shade constructions on his fields. We are particularly grateful for Stef den Hond and his continuous physical and mental support throughout the experiment. We also thank Marijn van de Broek, Laura Stefan, Anja Schmutz, Roman Hüppi, Simon Köldorfer, Johannes Blacha, Anne-Marlen Riemensperger, Christina Lagger, Simone Gerber, Lea Kreiselmeyer and Loes den Hond for their help during field work and Sabine Frick, Sophie Orivel, Anita Vaccaro and Michael Zens for their help after harvest. Furthermore, we thank UFA Samen and Sativa Rheinau for the provision of seeds, Landor for their consultation and provision of the biological fertilisers and Hortima for the provision of material for the shade net constructions.

## Notes

### Competing Interest Statement

The authors have declared no competing interest.

